# Toward an objective diagnostic for detection of *Leishmania RNA virus* (LRV) in clinical samples from cutaneous leishmaniasis patients

**DOI:** 10.1101/338095

**Authors:** Marcela Parra-Muñoz, Samanda Aponte, Clemencia Ovalle-Bracho, Carlos H. Saavedra, María C. Echeverry

## Abstract

*Leishmania RNA virus* (LRV) is a double strand RNA virus belonging to the *Totiviridae* family detected as cytoplasmic inclusions in some strains of the human parasite *Leishmania spp*. Experimental evidence supports the hypothesis that human co-infection with *Leishmania spp*-LRV triggers an exacerbated immune response in the host that can be responsible for the observed complicated outcomes in cutaneous leishmaniasis (CL), such as mucosal leishmaniasis (ML) and treatment failure of cutaneous leishmaniasis (TFCL). However, the reported frequencies of LRV associated to complicated outcomes in patients’ series are highly variable, diminishing the relevance on the virus presence in the pathogenesis of the disease. For determining if the apparent inconsistent information about the frequency of LRV associated to CL complicated outcomes could be connected with the virus detection approach, this study tested previously described methods for LRV detection in clinical samples of patients according with the type of sample. In 36 samples with diagnosis of complicated forms of CL (15 ML, 21 TFCL) and 6 samples with non-*Leishmania spp* infection, LRV presence was assessed by RT-PCR, RT-qPCR and nested-PCR.

By combining methods, LRV1 presence was confirmed in 45% (9/20) of isolates and 75% (12/16) of biopsies. The predominant LRV1-infected *Leishmania* species in this study was *Leishmania (Viannia) braziliensis* and, for first time, *Leishmania (Viannia) panamensis* was found infected in clinical samples. In a number of cases, LRV1 was undetectable in the isolates but present in their respective biopsies, suggesting the possibility of underreporting of LRV1 presence in studies focused in parasites isolates exclusively.

## Introduction

*Leishmania* species belonging to the *Viannia* subgenus are predominant in South America where they generate complicated forms of the disease such as mucosal leishmaniasis (ML) (1, 2) and treatment failure of cutaneous leishmaniasis (TFCL) (3, 4).

The occurrence of the above-mentioned complications has been associated with the presence of a cytoplasmic virus in the infecting parasite, known as *Leishmania RNA virus* (LRV). This virus belongs to the *Totiviridae* family, a group of RNA viruses present in other protozoa and fungi (5–9). Due to differences in sequence and genome organisation between LRVs associated to Old World and New World *Leishmania* species, they are classified as LRV2 and LRV1, respectively (10–13).

It has been demonstrated that metastasising (but not non-metastasising) strains of *Leishmania (Viannia) guyanensis* have high LRV1 burdens. In addition, there is experimental evidence that LRV presence induces hyper-immune responses to *Leishmania* infection inducing pro-inflammatory responses with high levels of TNFα, IL-6, CCL5 and CXCL10, similar to the immunological profile observed in patients suffering ML (14–16). Converging with this line of evidence is the observation that pro-inflammatory IL-17 levels are high in patients with chronic cutaneous leishmaniasis produced by *L. (V.) guyanensis-LRV1+* (17).

Supporting the hypothesis that LRV-*Leishmania spp* co-infection worsens disease prognosis through a Type 1 interferon response, murine model studies have shown that host co-infection with *L. (V.) guyanensis-LRV1+* or *L. (V.) guyanensis* plus exogenous interferon-inducing viruses (e.g., lymphocytic choriomeningitis virus or Toscana virus) produce similar clinical disease, in which relapse risk is increased secondary to parasite reactivation (18).

One clinical study has shown a strong association between LRV1 co-infection of *L. (V.) braziliensis* and pentavalent antimony treatment failure of CL, ML, and mucocutaneous leishmaniasis (19). Furthermore, the risk of CL relapse appears to increase during pentamidine treatment when *L. (V.) guyanensis* is infected with LRV1 (20).

At the epidemiologic level, LRV1/2 has been found at frequencies ranging from 0 to 87% in clinical samples from patients with leishmaniasis (15, 17, 19–33); the strength of the association between viral presence and complication development varies according to the study (24, 28) and among different South-American Regions (30). The wide range in LRV1 frequency of detection in clinical samples may reflect diverse experimental approaches that do not account for differences in clinical specimens.

To overcome detection biases in identifying LRV1 in clinical samples, the present study was designed to determine LRV1 frequencies in samples from complicated Colombian patients, suffering from TFCL or ML, through the use of complementary approaches taking into account the sample source, the parasite species and the viral load.

## Materials and Methods

### Ethics Statement

Research in this study was subject to ethical review and approved by the ethics committees from the participant institutions, in accordance with National (resolution 008430 of Colombian Health Ministry) and International (Declaration of Helsinki and amendments, World Medical Association, Korea 2008) guidelines. All clinical samples had been taken from patients as part of normal diagnosis and treatment, with no unnecessary invasive procedures, and with written informed consent. Guiding Principles for Biomedical Research Involving Animals (Council for International Organizations of Medical Sciences) were followed regarding animal experimentation.

### Type of study and Samples

The present study corresponds to a descriptive study with an experimental component using clinical samples (n=42) that had been collected previously. 21 samples collected from patients previously diagnosed as NRCL, of which 7 correspond to frozen biopsies and 14 to parasite isolates (34). 15 samples from patients with diagnosis of ML, of which 9 to frozen biopsies and 6 correspond to parasite isolates. 6 samples of frozen nasal biopsies from patients with confirmed diagnosis different to ML such as: lepromatous leprosy, traumatic piercing, sporotrichosis, chronic hyperplastic eosinophilic sinusitis, deep mycosis and squamous cell carcinoma, were included as negative controls.

### Parasite Isolates

Stocks of cryopreserved parasite isolates were thawed and seeded in Senekjie medium (35). Once adequate growth of the parasites was achieved, they were amplified until reaching the stationary phase in Schneider medium 10% FBS, at a temperature of 26 °C. Reference strains MHOM/BR/75/M4147 and MHOM/PA/71/LS94, grown on Schneider medium 10% FBS, were used as positive and negative controls for LRV1 infection respectively.

### Clinical samples and hamster biopsies

Fragments of about 3mm from patient’s biopsies were kept in dry sterile tube at -80 °C until RNA and DNA extraction. In order to standardise a protocol for LRV1 detection in tissue from biopsies, two young female golden hamsters (*Mesocricetus auratus*) were inoculated subcutaneously in footpads with 2×10^6^ MHOM/BR/75/M4147 metacyclic promastigotes. Three weeks after inoculation the animals were sacrificed and excisional biopsy were collected for proceeding to acid nucleic extraction.

### RNA extraction and Retrotranscription

Direct-zol™ RNA MiniPrep kit (Zymo Research) was used for the extraction RNA from 3.5×10^7^ to 3.4×10^8^ stationary phase promastigotes. For the biopsies, extraction was made using the Quick-RNA^TM^ MiniPrep Plus kit (Zymo Research) following the manufacturer’s instructions. cDNA was synthesised using High Capacity cDNA Reverse Transcription Kit (Thermo Fisher Scientific) following the manufacturer’s recommendations.

### Primers and 18S-*Leishmania spp* PCR amplification

Modified RT-PCR for *Leishmania spp*-18S following the protocol described by van den Bogaart (36) was performed. The set of primers used correspond to: Fw 5’- CCAAAGTGTGGAGATCGAAG -3’ and Rev 5’- GGCCGGTAAAGGCCGAATAG -3’.

### LRV1 Detection in cDNA from *Leishmania spp* isolates

The set of primers used for RT-PCR correspond to those described by Zangger et al (26); the ones used for RT-qPCR correspond to the set designed by Masayuki et al (27).

### LRV1 Detection in cDNA from biopsies and sequencing of nested-PCR products

The published nested-PCR protocol (24) was optimised by the use of cDNA from the *L. (V.) guyanensis*-LRV+ infected tissue from hamsters. The thermal profile included a cycle of 95 °C for 3 min; 35 cycles of 95 °C for 30 sec, 55 °C for 30 sec and 72 °C for 30 sec. The mixture was made to a final volume of 50 μl containing 1X Buffer KCl, 1.5 mM MgCl2, 200 μM dATP, dCTP, dTTP and dGTP, 2.5 units of recombinant Taq polymerase, 0.3 μM of each primer ANILRV1 (24): Fw 5’-CTGACTGGACGGGGGGTAAT-3’ and ANILRV1 Rev 5’- CAAAACACTCCCTTACGC-3’ were used in the first round and ANILRV-2 Fw 5’- GGTAATCGAGTGGGAGTCC-3’ and ANILRV-2 Rev 5’-GCGGCAGTAACCTGG-3’ in the second round. The amplified fragments were 125 bp and 90 bp for first and second round respectively and they were visualised in agarose gels. As a positive control cDNA from *L. (V.) guyanesis M4147* was used; negative controls correspond to cDNA from the 6 samples of frozen nasal biopsies from patients with confirmed diagnosis different to ML.

Fragments of 90 bp were purified from 1.5% agarose gels using the QIAquick gel extraction (Qiagen) and amplified products were delivered for sequencing to Macrogen Inc., Korea. The sequences obtained were analysed using the free Windows Software Molecular Evolutionary Genetics Analysis “MEGA 7”. Each consensus sequence was subjected to an identity test with NCBI’s Blast program.

#### Viral load quantification

A single plasmid standard was constructed containing a target sequence of 420pb corresponding to positions 16 to 435 in the 5’UTR region of LRV1. The sequence was amplified from MHOM/BR/75/M4147 cDNA using the previously described forward primer 5’ATGCCTAAGAGTTTGGATTCG-3’ (27) and a new designed reverse primer 5’-GAATTCCACGTGAACATACCTTGAGTATAC-3’. The PCR product was inserted into the PCR-4-TOPO^®^ TA Vector (K457502), following the manufacturer’s instructions and the construct amplified in *E. coli* Strain One Shot™ TOP10. To test the linearity of the single construct plasmid assay, standard curves of qPCR were created ranging from 2.3×10^1^ to 2.3×10^8^ plasmid copies per reaction. The qPCR was performed under previously described conditions and the observed Ct average for each sample was used to calculate its viral load by using the linear regression equation from the standard curve (Log10.ge vs Ct value).

### Leishmania species identification

Leishmania-species identification was made using hsp70 PCR-RFLP; species identification from isolates was performed according to Garcia and Montalvo et al. (37, 38) and for biopsies according to Cruz-Barrera et al. (39).

## Results

A multi-step process was used to create an LRV1 detection algorithm based on sample type. First, samples were classified as isolates (n=20) or biopsies (n= 16). In cases of isolates, RT-PCR (25, 26) was performed on cDNA from 19 *L. (V.) braziliensis* and one *L. (V.) panamensis*, isolates from lesions of patients with ML (n = 6) or TFCL (n=14). The expected products, ~485 bp corresponding to a viral genome region located at position 1089 to 1574, were observed in 9 out to 20 isolates (Fig. 1A).

**Figure 1.**
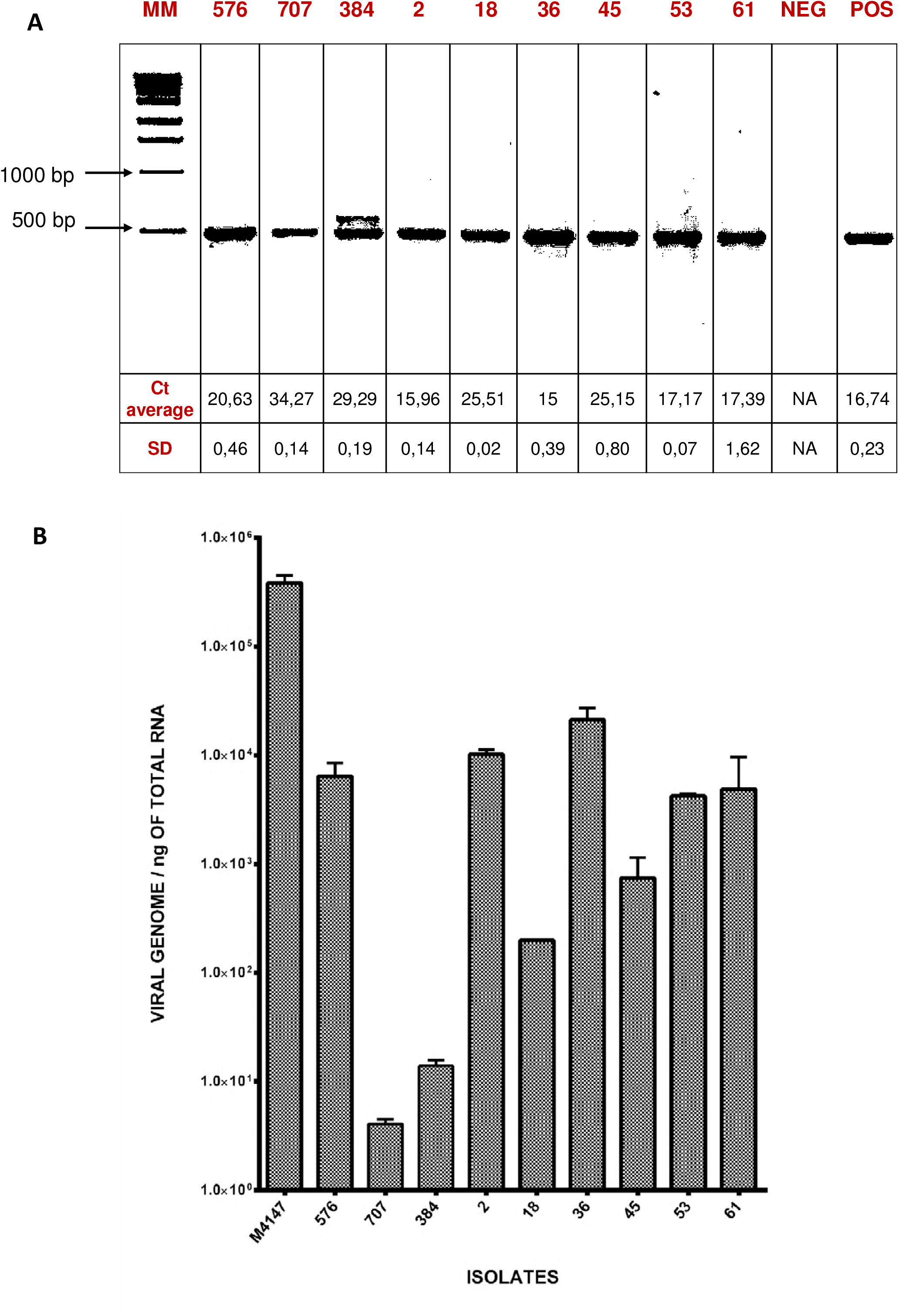
RT-PCR is useful for detection of low viral loads in *Leishmania spp*. isolates infected with LRV1. **A.** Amplification of LRV RNA capsid fragment (~ 488bp) using RT-PCR (26). Values on gel bottom correspond to the observed Ct for LRV1 5-’UTR amplification from the same isolates when detection was made by RT-qPCR. **B** The observed Cts displayed in A were used to calculate the viral loads of the samples. Numbers on top of the gel in A. and on the X axis in B. correspond to isolates identities.

Considering that different *Leishmania* strains can carry different LRV1 loads, an alternative detection method was used to determine whether a more conserved nucleotide sequence region could be used to improve detection sensitivity (10, 11, 13, 33). This detection was accomplished using a modified version of the RT-qPCR method described by *Massayuki et al*. (27), in which primers target the LRV 5’-UTR at position 16 to 239. This RT-qPCR of LRV 5’-UTR was used also to estimate the viral load in the isolates. RT-qPCR results were consistent with RT-PCR, detecting LRV1 presence in the same 9 isolates, even at low viral loads such as 4,09×10^−1^ viral genome/ng of total RNA (Fig. 1B and Table 1.).

**Table 1.**
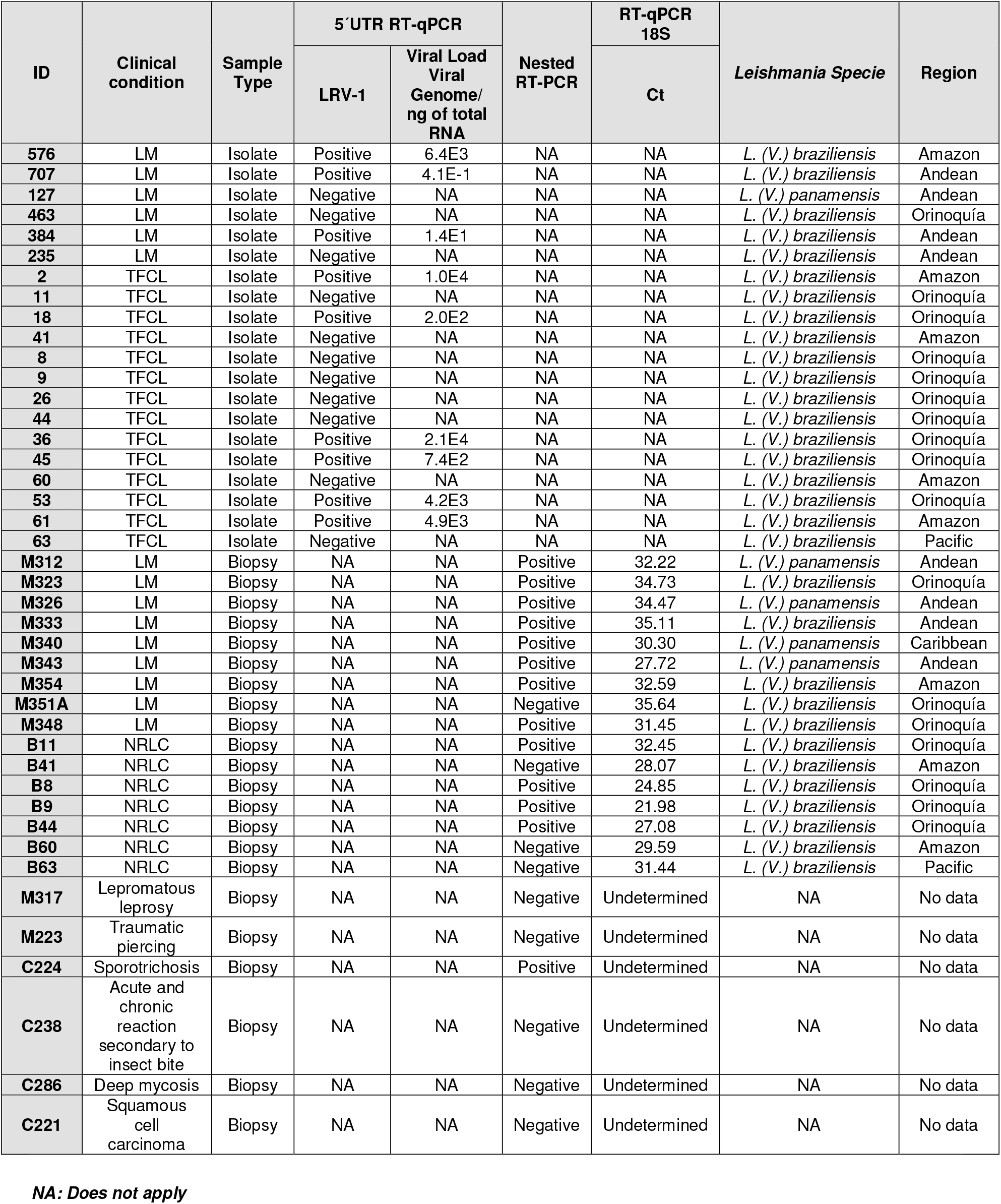
Features of the clinical samples included in this study.

| ID | Clinical condition | Sample Type | 5'UTR RT-qPCR |  | Nested RT-PCR | RT-qPCR 18S | *Leishmania Specie* | Region |
| --- | --- | --- | --- | --- | --- | --- | --- | --- |
|  |  |  | LRV-1 | Viral Load Viral Genome/ ng of total RNA |  | Ct |  |  |
| 576 | LM | Isolate | Positive | 6.4E3 | NA | NA | *L. (V.) braziliensis* | Amazon |
| 707 | LM | Isolate | Positive | 4.1E-1 | NA | NA | *L. (V.) braziliensis* | Andean |
| 127 | LM | Isolate | Negative | NA | NA | NA | *L. (V.) panamensis* | Andean |
| 463 | LM | Isolate | Negative | NA | NA | NA | *L. (V.) braziliensis* | Orinoquía |
| 384 | LM | Isolate | Positive | 1.4E1 | NA | NA | *L. (V.) braziliensis* | Andean |
| 235 | LM | Isolate | Negative | NA | NA | NA | *L. (V.) braziliensis* | Andean |
| 2 | TFCL | Isolate | Positive | 1.0E4 | NA | NA | *L. (V.) braziliensis* | Amazon |
| 11 | TFCL | Isolate | Negative | NA | NA | NA | *L. (V.) braziliensis* | Orinoquía |
| 18 | TFCL | Isolate | Positive | 2.0E2 | NA | NA | *L. (V.) braziliensis* | Orinoquía |
| 41 | TFCL | Isolate | Negative | NA | NA | NA | *L. (V.) braziliensis* | Amazon |
| 8 | TFCL | Isolate | Negative | NA | NA | NA | *L. (V.) braziliensis* | Orinoquía |
| 9 | TFCL | Isolate | Negative | NA | NA | NA | *L. (V.) braziliensis* | Orinoquía |
| 26 | TFCL | Isolate | Negative | NA | NA | NA | *L. (V.) braziliensis* | Orinoquía |
| 44 | TFCL | Isolate | Negative | NA | NA | NA | *L. (V.) braziliensis* | Orinoquía |
| 36 | TFCL | Isolate | Positive | 2.1E4 | NA | NA | *L. (V.) braziliensis* | Orinoquía |
| 45 | TFCL | Isolate | Positive | 7.4E2 | NA | NA | *L. (V.) braziliensis* | Orinoquía |
| 60 | TFCL | Isolate | Negative | NA | NA | NA | *L. (V.) braziliensis* | Amazon |
| 53 | TFCL | Isolate | Positive | 4.2E3 | NA | NA | *L. (V.) braziliensis* | Orinoquía |
| 61 | TFCL | Isolate | Positive | 4.9E3 | NA | NA | *L. (V.) braziliensis* | Amazon |
| 63 | TFCL | Isolate | Negative | NA | NA | NA | *L. (V.) braziliensis* | Pacific |
| M312 | LM | Biopsy | NA | NA | Positive | 32.22 | *L. (V.) panamensis* | Andean |
| M323 | LM | Biopsy | NA | NA | Positive | 34.73 | *L. (V.) braziliensis* | Orinoquía |
| M326 | LM | Biopsy | NA | NA | Positive | 34.47 | *L. (V.) panamensis* | Andean |
| M333 | LM | Biopsy | NA | NA | Positive | 35.11 | *L. (V.) braziliensis* | Andean |
| M340 | LM | Biopsy | NA | NA | Positive | 30.30 | *L. (V.) panamensis* | Caribbean |
| M343 | LM | Biopsy | NA | NA | Positive | 27.72 | *L. (V.) panamensis* | Andean |
| M354 | LM | Biopsy | NA | NA | Positive | 32.59 | *L. (V.) braziliensis* | Amazon |
| M351A | LM | Biopsy | NA | NA | Negative | 35.64 | *L. (V.) braziliensis* | Orinoquía |
| M348 | LM | Biopsy | NA | NA | Positive | 31.45 | *L. (V.) braziliensis* | Orinoquía |
| B11 | NRLC | Biopsy | NA | NA | Positive | 32.45 | *L. (V.) braziliensis* | Orinoquía |
| B41 | NRLC | Biopsy | NA | NA | Negative | 28.07 | *L. (V.) braziliensis* | Amazon |
| B8 | NRLC | Biopsy | NA | NA | Positive | 24.85 | *L. (V.) braziliensis* | Orinoquía |
| B9 | NRLC | Biopsy | NA | NA | Positive | 21.98 | *L. (V.) braziliensis* | Orinoquía |
| B44 | NRLC | Biopsy | NA | NA | Positive | 27.08 | *L. (V.) braziliensis* | Orinoquía |
| B60 | NRLC | Biopsy | NA | NA | Negative | 29.59 | *L. (V.) braziliensis* | Amazon |
| B63 | NRLC | Biopsy | NA | NA | Negative | 31.44 | *L. (V.) braziliensis* | Pacific |
| M317 | Lepromatous leprosy | Biopsy | NA | NA | Negative | Undetermined | NA | No data |
| M223 | Traumatic piercing | Biopsy | NA | NA | Negative | Undetermined | NA | No data |
| C224 | Sporotrichosis | Biopsy | NA | NA | Positive | Undetermined | NA | No data |
| C238 | Acute and chronic reaction secondary to insect bite | Biopsy | NA | NA | Negative | Undetermined | NA | No data |
| C286 | Deep mycosis | Biopsy | NA | NA | Negative | Undetermined | NA | No data |
| C221 | Squamous cell carcinoma | Biopsy | NA | NA | Negative | Undetermined | NA | No data |

**NA: Does not apply**

Previously to process samples corresponding to biopsies for LRV1 detection, qPCR for *Leishmania spp* 18S gene (36) was performed for all cDNA extracted from biopsy specimens (Table 1). This step allowed *Leishmania* infection to be ruled out in one of the biopsies diagnosed as LM and subsequently LRV1 detection efforts focused on specimens with detectable parasites avoiding a false-negative bias for LRV1 detection. qPCR for *Leishmania spp* 18S was also useful in order to provide evidence of the absence of parasites in all of the 6 specimens used for differential diagnosis and considered true negative controls for detection of LRV1 in biopsies (Table1).

A different approach was needed to detect LRV1 infections in biopsies given that PCR and qPCR methods were not feasible for LRV1 detection in an experimentally positive control consistent of a biopsy specimen from a (LVR+)-MHOM/BR/75/M4147 inoculated golden hamster (see Methods). Therefore, a nested PCR was used (24) for such purpose.

Nested PCR for LRV1 detection has been attempted with varying levels of success (24, 32). In this study, the protocol described by Ramos (24) was used after determining that the estimated amount of cDNA from a positive control biopsy could range from 6.6 to 266 ng without inhibiting detection. Following nested PCR standardisation and for subsequent clinical sample analyses, cDNA from (LVR+)-MHOM/BR/75/M414 was used as a positive control and cDNAs from 6 nasal mucosa biopsies from patients with non-*Leishmania spp* (proven differential diagnosis) (Table 1) were used as a negative control. The resulting products had a single band of 90 bp, which was detected independently of the cDNA amount.

The nested RT-PCR technique was applied to 22 biopsies specimens from patients with ML (n =9), TFCL (n = 7) and with proven different (non-ML) diagnosis (n = 6). The virus was detected in eight out of nine ML specimens and four out of seven TFCL specimens. One nasal biopsy specimen diagnosed as sporotrichosis also yielded a positive result for amplification of a fragment of 90bp corresponding to position 167-254 of the LRV1 5’-UTR. Sequencing data from the PCR products showed that in all cases the sequence obtained correspond to LRV1 5’-UTR with an identity ranging from 91-100% with other leishmaniavirus sequences.

The highest frequency of LRV1-positivity was observed in biopsy specimens, and the lowest frequency was observed in isolates (Table 1). Considering that seven patients with TFCL contributed both biopsy specimens and isolate samples, an analysis was conducted to simultaneously evaluate the LRV1 presence in biopsies and in their respective isolates (Table 1: B11, B41, B8, B9, B44, B60, and B63 and 11, 41, 8, 9, 44, 60, and 6). LRV1 presence was established in four biopsy samples in which their respective isolates showed no LRV1 presence assessed by RT-PCR and RT-qPCR (Table 1: B8 vs 8, B9 vs 9, B11 vs 11 and B44 vs 44).

## Discussion

Colombia has one of the world’s highest cutaneous leishmaniasis (CL) incidences (40, 41). Approximately 10,000 new cases are reported each year, with *L. (V) panamensis* and *L. (V) braziliensis* being the most common agents associated to human disease (42–44). The complication rates for ML and TFCL are not well known but have been estimated in other South American regions. Past studies report that 10% of CL cases associated with *L. (V) braziliensis* result in ML (42, 43), and 25% of CL cases results in TFCL (45, 46). Hence, a more detailed epidemiologic picture is urgently needed to better understand CL prognosis. Understanding the role of LRV1 in human CL outcomes requires an ambitious epidemiologic design and robust detection methods using a multicentre study. Therefore, we aimed to evaluate the possible biases in LRV1 detection when using the simplest and cheapest available approaches (24–27).

Our use of complementary approaches in this study led us to the important finding that LVR1 is apparently lost during *in vitro* parasite growth. An explanation for the biological events responsible for this loss was not addressed in the present study. However, it is important to note that if this viral loss commonly occurs among other *Leishmania* strains, previous studies using LRV detection exclusively in isolates may have underestimated the rate of viral co-infection for at least some *Leishmania* species (17, 19–21, 25, 29–32).

Other factors involved in poor detection of LRV by PCR or qPCR in clinical samples are technical conditions linked to test performance. When comparing PCR for LRV1 detection using “universal primers” (26) with the viral loads, it was observed that weaker signals correspond to samples with lower viral loads (Figure 1). It is important to note that observed low viral loads, as scarce as 4.09*10^−1^ viral genome/ng of RNA seems to indicate that not all the parasites are infected during the *in vitro* growth; however, this low viral presence was detectable by a simple RT-PCR (26). In the present study, before ruling out an isolate as LRV1-negative, multiple attempts for detection were carried out with variable amounts of RNA from 6 to 200 ng (data not shown).

Here, LRV1 detection frequency in specimens from patients with ML was higher when using biopsy specimens with nested PCR (88.9%) than when using isolates (50%), this finding is consistent with other studies showing that direct sample analysis results in higher rates of LRV1 detection in association with ML (27, 28). However, other authors who simultaneously analysed clinical samples from patients with CL and ML found that LRV1 is not preferentially associated with ML (19). This discrepancy may be explained by two factors. First, the previous study used isolates (not biopsy specimens) to assess LRV presence. Second, ML is a long-term complication (47, 48), and it is not possible to determine in mixed patient populations what percent of those with CL will eventually develop ML.

In samples derived from patients with TFCL, LRV1 was also detected more frequently among biopsy specimens than among isolates (57.14% vs 42.86%). The predominant species analysed in this study was *L. (V) braziliensis;* however, four ML cases were associated with *L.(V) panamensis*.

Although there is a geographic bias in this study and a reduced number of samples, with only one sample from the Pacific region and one sample from the Caribbean region, the LRV1+ frequency for the remaining Colombian regions ranged from 57% to 69%. Thus, *Leishmania* strains harbouring LRV1 may have spread since 1996, when a report of 69 *Leishmania* strains showed low LRV1 parasite infection rates, found exclusively in the Amazon region (21). Albeit that previous report included 26 *L. (V) panamensis* strains, none were LRV1-infected. In contrast, four out of five biopsy specimens from patients in this study with ML associated with *L. (V) panamensis* were positive for LRV1 presence (Table 1). This finding is not only relevant with respect to the LRV1 distribution outside of the Amazon region, but also because this is the first report of LRV1 infection of this parasite species in Colombia, where the predominant species associated to human infections is *L. (V) panamensis*. An alternative explanation for differences between the present study and the aforementioned study is that the previous study used only parasites isolates, whereas our *L. (V) panamensis*-LRV1+ results were found in biopsy specimens from patients with complicated CL.

Recently, the presence of LRV1 in *L. (V) panamensis* isolates from Ecuador and Costa Rica was reported showing that LRV1 is broadly spread into *Leishmania* species from the *Viannia* subgenus at the north of the Amazon Region (15).

Finally, a noteworthy observation that requires further investigation is the detection of LRV1 in a patient with nasal sporotrichosis with no evidence of *Leishmania* infection as judged from 18S qPCR detection. This result, provides further evidence of viruses from the *Totiviradae* family infecting fungi of Ophiostoma family like *Ophiostoma minus* (49), and *Sporothrix schenckii* (50). The amplified sequence derived from the nested RT-PCR of this sample showed a 91-100% percentage of similarity with the LVR1 region comprising nucleotides 167-256 derived from various Leishmania isolates; however, it is worth noting the reduced size of this amplification product (90bp). This preliminary evidence is in accordance with other authors data that, when comparing the available genomic regions of *Ophiostoma minus totivirus* and other mycoviruses belonging to the *Totiviridae* family, the analysis shows that they do not differ highly from viruses sampled from non-fungal hosts such as *Leishmania RNA virus* (51).

As a conclusion, findings in the present study suggest that detection of LRV1 in epidemiological studies aimed at determining an association between LRV1 presence and complicated outcomes of CL, should consider the evaluation of LRV1 occurrence using samples coming directly from the patients such as biopsies of lesion swaps. Simple and cheap methods such as nested PCR and RT-PCR could be useful for that purpose.

## Acknowledgments

Authors render thanks to Dr. Stephen Beverley for his help in improving the detection methods and data discussion and Dr. Jussep Salgado for helping with *Leishmania* species identification. This work was supported by Universidad Nacional de Colombia grant HERMES-32319& 28542. Marcela Parra-Muñoz was funded by Colciencias young researchers programme.

## References

1. Strazzulla A, Cocuzza S, Pinzone MR, Postorino MC, Cosentino S, Serra A, Cacopardo B, Nunnari G. 2013. Mucosal leishmaniasis: an underestimated presentation of a neglected disease. Biomed Res Int 2013:1–7.

2. de Oliveira Guerra JA, Prestes SR, Silveira H, Câmara LIdAR, Gama P, Moura A, Amato V, Barbosa MdGV, de Lima Ferreira LC. 2011. Mucosal leishmaniasis caused by Leishmania (Viannia) braziliensis and Leishmania (Viannia) guyanensis in the Brazilian Amazon. PLoS Negl Trop Dis 5:1–5.

3. Haldar AK, Sen P, Roy S. 2011. Use of antimony in the treatment of leishmaniasis: current status and future directions. Mol Biol Int 2011: 1–23.

4. Vanaerschot M, Dumetz F, Roy S, Ponte-Sucre A, Arevalo J, Dujardin J-C. 2014. Treatment failure in leishmaniasis: drug-resistance or another (epi-) phenotype. Expert Rev Anti Infect Ther 12:937–946.

5. Ghabrial SA. 2008. Totiviruses, p163–174. *In* Regenmortel, Brian W. J. MahyMarc H. V. Van (ed), Encyclopedia of Virology (Third Edition). Academic Press, Oxford.

6. Diamond LS, Mattern CFT. 1976. Protozoal Viruses, p87–112. *In* Max A. Lauffer, Frederik B. Bang, Karl Maramorosch, Kenneth M. Smith (ed), Advances in Virus Research, vol 20. Academic Press.

7. Akopyants NS, Lye L-F, Dobson DE, Lukeš J, Beverley SM. 2016. A novel bunyavirus-like virus of trypanosomatid protist parasites. Genome Announc 4:1–2.

8. Mertens P. 2004. The dsRNA viruses. Virus Res 101:3–13.

9. Wang A, Wang C. 1991. Viruses of the protozoa. Annu Rev Microbiol 45:251–263.

10. Stuart KD, Weeks R, Guilbride L, Myler PJ. 1992. Molecular organization of Leishmania RNA virus 1. Proc Natl Acad Sci U S A 89:8596–8600.

11. Guilbride L, Myler PJ, Stuart K. 1992. Distribution and sequence divergence of LRV1 viruses among different Leishmania species. Mol Biochem Parasitol 54:101–104.

12. Scheffter SM, Ro YT, Chung IK, Patterson J. 1995. The complete sequence of Leishmania RNA virus LRV2–1, a virus of an Old World parasite strain. Virology 212:84–90.

13. Scheffter S, Widmer G, Patterson JL. 1994. Complete sequence of Leishmania RNA virus 1–4 and identification of conserved sequences. Virology 199:479–483.

14. Ives A, Ronet C, Prevel F, Ruzzante G, Fuertes-Marraco S, Schutz F, Zangger H, Revaz-Breton M, Lye LF, Hickerson SM, Beverley SM, Acha-Orbea H, Launois P, Fasel N, Masina S. 2011. Leishmania RNA virus controls the severity of mucocutaneous leishmaniasis. Science 331:775–8.

15. Kariyawasam R, Grewal J, Lau R, Purssell A, Valencia BM, Llanos-Cuentas A, Boggild AK. 2017. Influence of Leishmania RNA Virus-1 on Pro-Inflammatory biomarker expression in a human macrophage model of American Tegumentary Leishmaniasis. J Infect Dis 216: 877–886.

16. Soliman MF, Ibrahim MM. 2017. Do parasite viruses affect the relationship of parasites with their host?. Edorium J Infect Dis 3:9–11.

17. Hartley M-A, Bourreau E, Rossi M, Castiglioni P, Eren RO, Prevel F, Couppié P, Hickerson SM, Launois P, Beverley SM. 2016. Leishmaniavirus-dependent metastatic leishmaniasis is prevented by blocking IL-17A. PLoS Pathog 12:1–19.

18. Rossi M, Castiglioni P, Hartley M-A, Eren RO, Prével F, Desponds C, Utzschneider DT, Zehn D, Cusi MG, Kuhlmann FM. 2017. Type I interferons induced by endogenous or exogenous viral infections promote metastasis and relapse of leishmaniasis. Proc Natl Acad Sci U S A 114:4987–92.

19. Adaui V, Lye L-F, Akopyants NS, Zimic M, Llanos-Cuentas A, Garcia L, Maes I, De Doncker S, Dobson DE, Arevalo J. 2015. Association of the endobiont double-stranded RNA virus LRV1 with treatment failure for human Leishmaniasis caused by Leishmania braziliensis in Peru and Bolivia. J Infect Dis 213:112–21.

20. Bourreau E, Marine Ginouves M, Prévot G. 2015. Leishmania-RNA virus presence in L. guyanensis parasites increases the risk of first-line treatment failure and symptomatic relapse. J Infect Dis 213:105–11.

21. Salinas G, Zamora M, Stuart K, Saravia N. 1996. Leishmania RNA viruses in Leishmania of the Viannia subgenus. Am J Trop Med Hyg 54:425–9.

22. Saiz M, Llanos-Cuentas A, Echevarria J, Roncal N, Cruz M, Muniz MT, Lucas C, Wirth D, Scheffter S, Magill A. 1998. Short report: detection of Leishmaniavirus in human biopsy samples of leishmaniasis from Peru. Am J Trop Med Hyg 58:192–4.

23. Ogg MM, Carrion R, Botelho ACDC, Mayrink W, Correa-Oliveira R, Patterson JL. 2003. Short report: quantification of leishmaniavirus RNA in clinical samples and its possible role in pathogenesis. Am J Trop Med Hyg 69:309–13.

24. Ramos Pereira LdO, Maretti-Mira AC, Rodrigues KM, Lima RB, Oliveira-Neto MPd, Cupolillo E, Pirmez C, Oliveira MPd. 2013. Severity of tegumentary leishmaniasis is not exclusively associated with Leishmania RNA virus 1 infection in Brazil. Mem Inst Oswaldo Cruz 108:665–667.

25. Zangger H, Ronet C, Desponds C, Kuhlmann FM, Robinson J, Hartley MA, Prevel F, Castiglioni P, Pratlong F, Bastien P, Muller N, Parmentier L, Saravia NG, Beverley SM, Fasel N. 2013. Detection of Leishmania RNA virus in Leishmania parasites. PLoS Negl Trop Dis 7:1–11.

26. Zangger H, Hailu A, Desponds C, Lye LF, Akopyants NS, Dobson DE, Ronet C, Ghalib H, Beverley SM, Fasel N. 2014. Leishmania aethiopica field isolates bearing an endosymbiontic dsRNA virus induce pro-inflammatory cytokine response. PLoS Negl Trop Dis 8:1–10.

27. Ito MM, Catanhêde LM, Katsuragawa TH, da Silva Junior CF, Camargo LMA, de Godoi Mattos R, Vilallobos-Salcedo JM. 2015. Correlation between presence of Leishmania RNA virus 1 and clinical characteristics of nasal mucosal leishmaniosis. Braz J Otorhinolaryngol 81:533–40.

28. Cantanhêde LM, da Silva Júnior CF, Ito MM, Felipin KP, Nicolete R, Salcedo JMV, Porrozzi R, Cupolillo E, Ferreira RdGM. 2015. Further evidence of an association between the presence of Leishmania RNA Virus 1 and the mucosal manifestations in Tegumentary Leishmaniasis patients. PLoS Negl Trop Dis 9:1–11.

29. Alves-Ferreira EVC, Toledo JS, De Oliveira AHC, Ferreira TR, Ruy PC, Pinzan CF, Santos RF, Boaventura V, Rojo D, López-Gonzálvez Á, Rosa JC, Barbas C, Barral-Netto M, Barral A, Cruz AK. 2015. Differential gene expression and infection profiles of cutaneous and mucosal Leishmania braziliensis isolates from the same patient. PLoS Negl Trop Dis 9:1–19.

30. Macedo DH, Menezes-Neto A, Rugani JM, Rocha AC, Silva SO, Melo MN, Lye L-F, Beverley SM, Gontijo CM, Soares RP. 2016. Low frequency of LRV1 in Leishmania braziliensis strains isolated from typical and atypical lesions in the State of Minas Gerais, Brazil. Mol Biochem Parasitol 210:50–4.

31. Ginouvès M, Simon S, Bourreau E, Lacoste V, Ronet C, Couppié P, Nacher M, Demar M, Prévot G. 2015. Prevalence and distribution of Leishmania RNA Virus 1 in Leishmania parasites from French Guiana. Am J Trop Med Hyg 94:102–6.

32. Hajjaran H, Mahdi M, Mohebali M, Samimi-Rad K, Ataei-Pirkooh A, Kazemi-Rad E, Naddaf SR, Raoofian R. 2016. Detection and molecular identification of Leishmania RNA virus (LRV) in Iranian Leishmania species. Arch Virol 161:3385–90.

33. Tirera S, Ginouves M, Donato D, Caballero IS, Bouchier C, Lavergne A, Bourreau E, Mosnier E, Vantilcke V, Couppié P. 2017. Unraveling the genetic diversity and phylogeny of Leishmania RNA virus 1 strains of infected Leishmania isolates circulating in French Guiana. PLoS Negl Trop Dis 11:1–20.

34. Perez-Franco JE, Cruz-Barrera ML, Robayo ML, Lopez MC, Daza CD, Bedoya A, Mariño ML, Saavedra CH, Echeverry MC. 2016. Clinical and parasitological features of patients with American Cutaneous Leishmaniasis that did not respond to treatment with Meglumine Antimoniate. PLoS Negl Trop Dis 10:1–13.

35. Walton BC, Shaw JJ, Lainson R. 1977. Observations on the in vitro cultivation of Leishmania braziliensis. J Parasitol 63:1118–19.

36. van den Bogaart E, Schoone GJ, Adams ER, Schallig HD. 2014. Duplex quantitative reverse-transcriptase PCR for simultaneous assessment of drug activity against Leishmania intracellular amastigotes and their host cells. Int J Parasitol Drugs Drug Resist 4:14–19.

37. Garcia L, Kindt A, Bermudez H, Llanos-Cuentas A, De Doncker S, Arevalo J, Tintaya KWQ, Dujardin J-C. 2004. Culture-independent species typing of neotropical Leishmania for clinical validation of a PCR-based assay targeting heat shock protein 70 genes. J Clin Microbiol 42:2294–97.

38. Montalvo AM, Fraga J, Monzote L, Montano I, De Doncker S, Dujardin JC, Van der Auwera G. 2010. Heat-shock protein 70 PCR-RFLP: a universal simple tool for Leishmania species discrimination in the New and Old World. Parasitology 137:1159–68.

39. Cruz Barrera M, Ovalle Bracho C, Ortegón Vergara V, Pérez Franco JE, Echeverry MC. 2015. Improving Leishmania Species Identification in Different Types of Samples from Cutaneous Lesions. J Clin Microbiol 53:1339–41.

40. World Health Organization. 2015. Status of endemicity of cutaneous leishmaniasis worldwine, 2015. Available at: http://www.who.int/leishmaniasis/burden/Status_of_endemicity_of_CL_worldwide_2015_with_imported_cases.pdf.

41. World Health Organization. 2009. Leishmaniasis: burden of disease. Available at: http://wwwwhoint/leishmaniasis/burden/en.

42. Ovalle CE, Porras L, Rey M, Ríos M, Camargo YC. 2006. Distribución geográfica de especies de Leishmania aisladas de pacientes consultantes al Instituto Nacional de Dermatología Federico Lleras Acosta, ESE, 1995–2005. Biomédica 26:145–51.

43. Corredor A, Kreutzer RD, Tesh RB, Boshell J, Palau MT, Caceres E, Duque S, Pelaez D, Rodriguez G, Nichols S. 1990. Distribution and etiology of Leishmaniasis in Colombia. Am J Trop Med Hyg 42:206–214.

44. Patino LH, Mendez C, Rodriguez O, Romero Y, Velandia D, Alvarado M, Pérez J, Duque MC, Ramírez JD. 2017. Spatial distribution, Leishmania species and clinical traits of Cutaneous Leishmaniasis cases in the Colombian army. PLoS Negl Trop Dis 11:1–15.

45. Llanos-Cuentas A, Tulliano G, Araujo-Castillo R, Miranda-Verastegui C, Santamaria-Castrellon G, Ramirez L, Lazo M, De Doncker S, Boelaert M, Robays J. 2008. Clinical and parasite species risk factors for pentavalent antimonial treatment failure in cutaneous leishmaniasis in Peru. Clin Infect Dis 46:223–231.

46. Tuon FF, Amato VS, Graf ME, Siqueira AM, Nicodemo AC, Neto VA. 2008. Treatment of New World cutaneous leishmaniasis-a systematic review with a meta-analysis. Int J Dermatol 47:109–124.

47. Amato VS, Tuon FF, Bacha HA, Neto VA, Nicodemo AC. 2008. Mucosal leishmaniasis: Current scenario and prospects for treatment. Acta Trop 105:1–9.

48. Osorio LE, Castillo CM, Ochoa MT. 1998. Mucosal leishmaniasis due to Leishmania (Viannia) panamensis in Colombia: clinical characteristics. Am J Trop Med Hyg 59:49–52.

49. Doherty M, Sanganee K, Kozlakidis Z, Coutts RH, Brasier C, Buck K. 2007. Molecular characterization of a totivirus and a partitivirus from the genus Ophiostoma. J Phytopatho 155:188–192.

50. Berbee ML, Taylor JW. 1992. 18S Ribosomal RNA gene sequence characters place the human pathogen Sporothrix schenckii in the genus Ophiostoma. Exp Mycol 16:87–91.

51. Göker M, Scheuner C, Klenk H-P, Stielow JB, Menzel W. 2011. Codivergence of mycoviruses with their hosts. PLoS One 6:1–13.

